# Sox2 modulation increases naïve pluripotency plasticity

**DOI:** 10.1101/2020.01.14.906933

**Authors:** Kathryn Tremble, Giuliano G. Stirparo, Lawrence E. Bates, Katsiaryna Maskalenka, Hannah T. Stuart, Kenneth Jones, Amanda Andersson-Rolf, Aliaksandra Radzisheuskaya, Bon-Kyoung Koo, Paul Bertone, José C. R. Silva

## Abstract

Induced pluripotency provides a tool to explore the molecular mechanisms underlying the establishment, maintenance and differentiation of naïve pluripotent stem cells (nPSCs). Here, we report that self-renewal of nPSCs requires minimal Sox2 expression (Sox2-low). Sox2-low nPSCs do not show impaired neuroectoderm specification and differentiate efficiently *in vitro* into all embryonic germ lineages. Strikingly, Sox2-low cells also differentiate towards the trophoblast lineage both *in vitro* and *in vivo*. At the single-cell level self-renewing Sox2-low nPSCs exhibit a homogeneous naïve molecular signature. However, they also display a basal trophoblast molecular signature and decreased ability of Oct4 to bind naïve-associated regulatory sequences compared to control cells. These features underlie observed enhanced cell potency upon the removal of self-renewing cues. In sum, this work defines Sox2 as a restrictor of developmental potential and suggests perturbation of the naïve pluripotent network as an underlying cause of increased cell potency.

**Highlights:**

- Low Sox2 expression is sufficient for naïve pluripotent stem cell self-renewal
- Low Sox2 expression does not impair neurectoderm differentiation *in vitro*
- Low Sox2 expression impairs Oct4 genomic occupancy
- Low Sox2 expression increases naïve pluripotent cell plasticity *in vitro* and *in vivo*

## Introduction

The naïve epiblast is pluripotent as it has the potential to differentiate into any cell type of the embryo proper but cannot form extraembryonic lineages. Naive Pluripotency can be captured *in vitro* from the epiblast in the form of embryonic stem cells (ESCs) (Evans and Kaufman, 1981; Martin, 1981) and through reprogramming of differentiated cells in the form of induced pluripotent stem cells (iPSCs) (Takahashi and Yamanaka, 2006), allowing us to study the molecular mechanisms underlying this identity. Recently it has been demonstrated that treatment of ESCs with specific small molecules induces expanded differentiation potential to include extraembryonic lineages (Yang et al., 2017a, 2017b). However, the mechanism underlying this has not been elucidated.

Sox2 is a member of the SRY-related HMG-box family of transcription factors (Wright et al., 1993) and is a core pluripotency factor. Sox2 knockout in embryos results in peri-implantation lethality and its deletion in ESCs results in loss of self-renewal with the cells becoming trophoblast-like stem cells (Avilion et al., 2003; Masui et al., 2007). Sox2 was originally discovered as a putative DNA-binding partner of the central pluripotency factor Oct4 (Chew et al., 2005; Rodda et al., 2005; Yuan et al., 1995). However, self-renewing *Sox2−/−* ESCs were generated in the presence of constitutive Oct4 expression, suggesting that the main role of Sox2 is to maintain Oct4 expression (Masui et al., 2007). Additionally, overexpression of Sox2 results in differentiation of ESCs (Kopp et al., 2008). It has been hypothesised that Sox2 acts as a neurectoderm specifier, which needs to be in balance with mesendoderm specifiers to result in pluripotency (Loh and Lim, 2011; Thomson et al., 2011). Therefore, the role and requirement of Sox2 in naïve pluripotency is currently unclear.

Here we utilised the process of induced pluripotency to investigate the biological role of Sox2. This uncovered the surprising role of Sox2 in restricting the potency of naïve pluripotent stem cells to embryonic lineages only. This impacts on both the understanding of the role of core naïve pluripotency factors and on the molecular basis governing the potency of naïve pluripotent stem cells (nPSCs).

## Results

### Low Sox2 expression is compatible with naïve pluripotent stem cell self-renewal

To explore the role of Sox2 in the process of induced pluripotency, we attempted to generate *Sox2−/−* iPSCs. *Sox2−/−* neural stem cells (NSCs) were generated using CRISPR/Cas9 (Figure S1A-D). Clonal lines were confirmed to have a frame-shift deletion in the Sox2 codon, resulting in loss of Sox2 protein (Figure S1A and B). The *Sox2−/−* NSCs maintained neural stem cell morphology, proliferative ability, and expression of neural stem cell markers (Figure S1C and D). The *Sox2−/−* NSCs contained GFP and the blasticidin resistance genes under the endogenous Rex1 regulatory sequences to allow identification and selection for naïve pluripotent stem cell identity (Wray et al., 2011). *Sox2−/−* NSCs were induced to reprogram by combining four or three of the classic Yamanaka retroviral factors: cMyc, Klf4, Oct4 and Sox2 (rMKOS) or cMyc, Klf4 and Oct4 (rMKO) (Takahashi and Yamanaka, 2006) (Figure 1A). Retroviral promoters are active in somatic cells but become epigenetically silenced in naïve pluripotency (Hotta and Ellis, 2008). Therefore we hypothesised that the retroviral Sox2 would drive reprogramming but become silenced after stabilisation of the network. We also used defined culture conditions containing inhibitors of Mek/Erk and Gsk3b signalling (2i) supplemented with LIF for optimal reprogramming efficiency (Silva et al., 2008). *Sox2−/−* NSCs were able to reprogram upon addition of a constitutive exogenous Sox2 transgene (Figure 1B). In the absence of retroviral Sox2, *Sox2−/−* NSCs failed to upregulate the Rex1-GFP reporter (Figure 1B). Surprisingly, multiple *Sox2−/−* Rex1-GFP+ colony-like groups of cells emerged when using rMKOS (Figure 1B). An independent experiment showed this to be 5 times less efficient compared to the reprogramming of WT NSCs (Figure 1C). Reprogrammed Sox2−/− rMKOS iPSCs were passagable in 2iLIF culture conditions and expressed naïve pluripotency markers (Figure 1D and 1E). Surprisingly, they also expressed Sox2 protein at very low levels (Figure 1F), which was due to a failure to fully silence retroviral Sox2 (Figure 1G). Importantly, this data indicates a strong selective pressure for cells expressing a minimal level of Sox2 protein and suggests that low Sox2 expression is compatible with maintenance of a naïve pluripotent stem cell molecular identity. Hereafter, we will refer to these *Sox2−/−* rMKOS iPSCs as Sox2-low iPSCs for simplicity.

**Figure 1.**
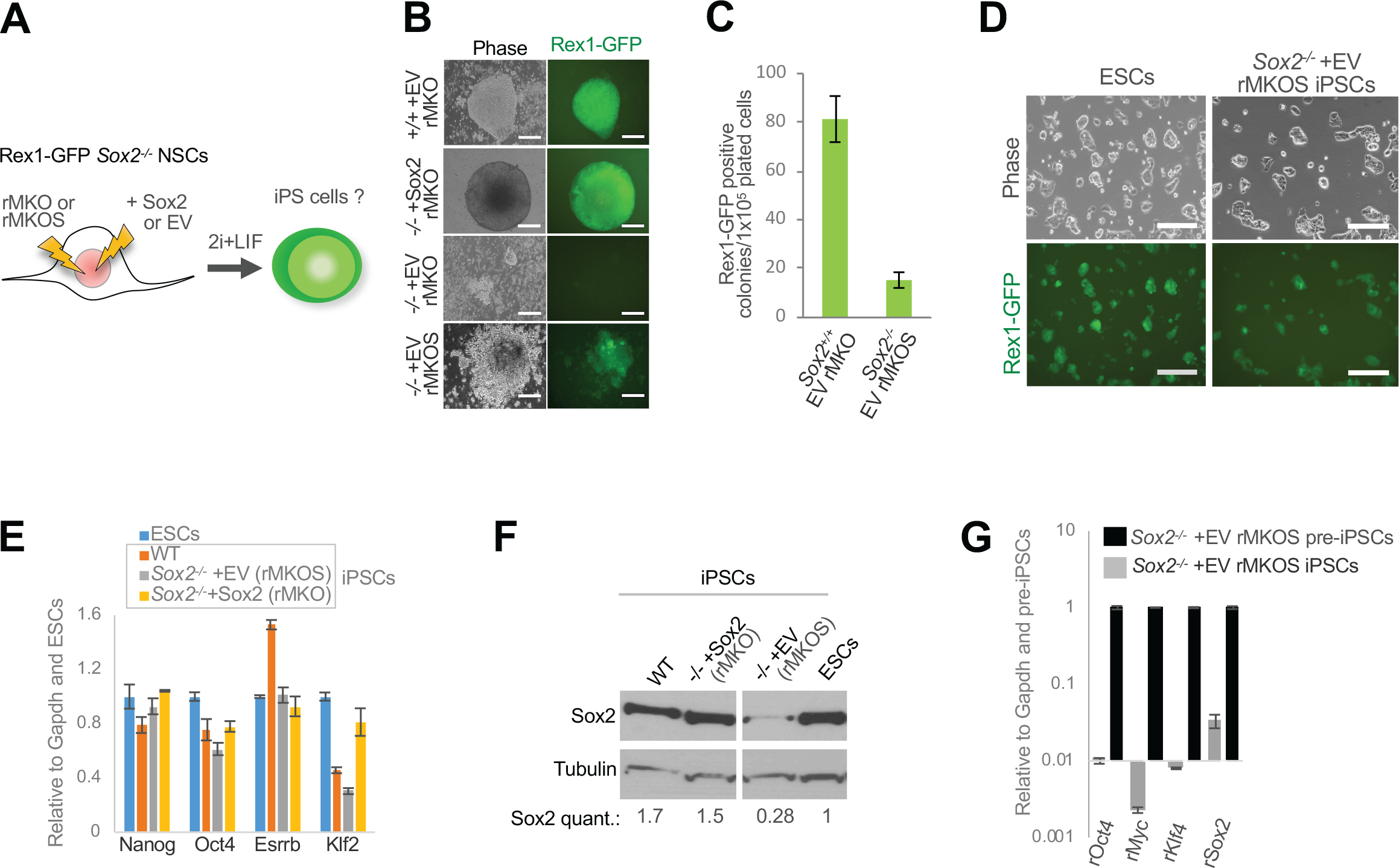
Generation of self-renewing iPSCs expressing low levels of Sox2. A) Diagram of experimental design. rMKO and rMKOS indicate use of retroviral (r) vectors containing reprogramming cMyc (M), Klf4 (K), Oct4 (O) and Sox2 (S) transgenes. + EV (empty vector) or + Sox2 represent use of a PiggyBac plasmid containing a CAG promoter driving constitutive expression of either an empty or Sox2 transgene respectively. B) Phase and GFP images of emerging Rex1-GFP+ iPSC colonies (n=3). C) Rex1-GFP+ colony counts for indicated genotypes (n=3). D) Phase and GFP images of Sox2−/− rMKOS iPSCs in 2iLIF post-selection. Rex1-GFP ESCs are shown as control. E) qRT-PCR analysis for indicated pluripotency associated factors in iPSC and control ESC lines. F) Western blot of Sox2 (≈40kDa) and Tubulin (≈50kDa) in iPSC and control ESC lines in 2iLIF with Sox2 quantification relative to ESCs, normalised to tubulin. Gap in Western blot represents removal of a non-relevant lane and image corresponds to same film exposure G) qRT-PCR of retroviral Sox2, Oct4, Klf4 and cMyc expression in Sox2−/− rMKOS iPSCs relative to preiPSCs. Scale bars = 200µm. Error bars indicate standard deviation of replicate qPCR reactions (n=3).

### Sox2-low iPSCs differentiate in serum plus LIF self-renewing conditions

It has previously been reported that Sox2 repression in ESCs results in loss of pluripotency and differentiation towards the trophoblast lineage in serum plus LIF (SLIF) conditions (Masui et al., 2007). Therefore, we attempted to culture Sox2-low iPSCs in SLIF. Strikingly, Sox2-low iPSCs downregulated the Rex1-GFP reporter, and some cells gained a trophoblast-like morphology (Figure 2A). They also downregulated pluripotency marker expression and upregulated trophoblast markers (Figure 2B-C). In addition, cells were not passageable, demonstrating loss of self-renewal. To ensure that the differentiation phenotype was due to low Sox2 expression, we generated Sox2-low rescue iPSCs by transfecting these with a constitutive Sox2 transgene (CAG-Sox2) in 2iLIF (Figure 2D). Upon SLIF medium switch, and in contrast to Sox2-low iPSCs, rescue iPSCs self-renewed, maintained naïve pluripotent gene expression and did not upregulate trophoblast marker expression (Figure 2E-F).

**Figure 2.**
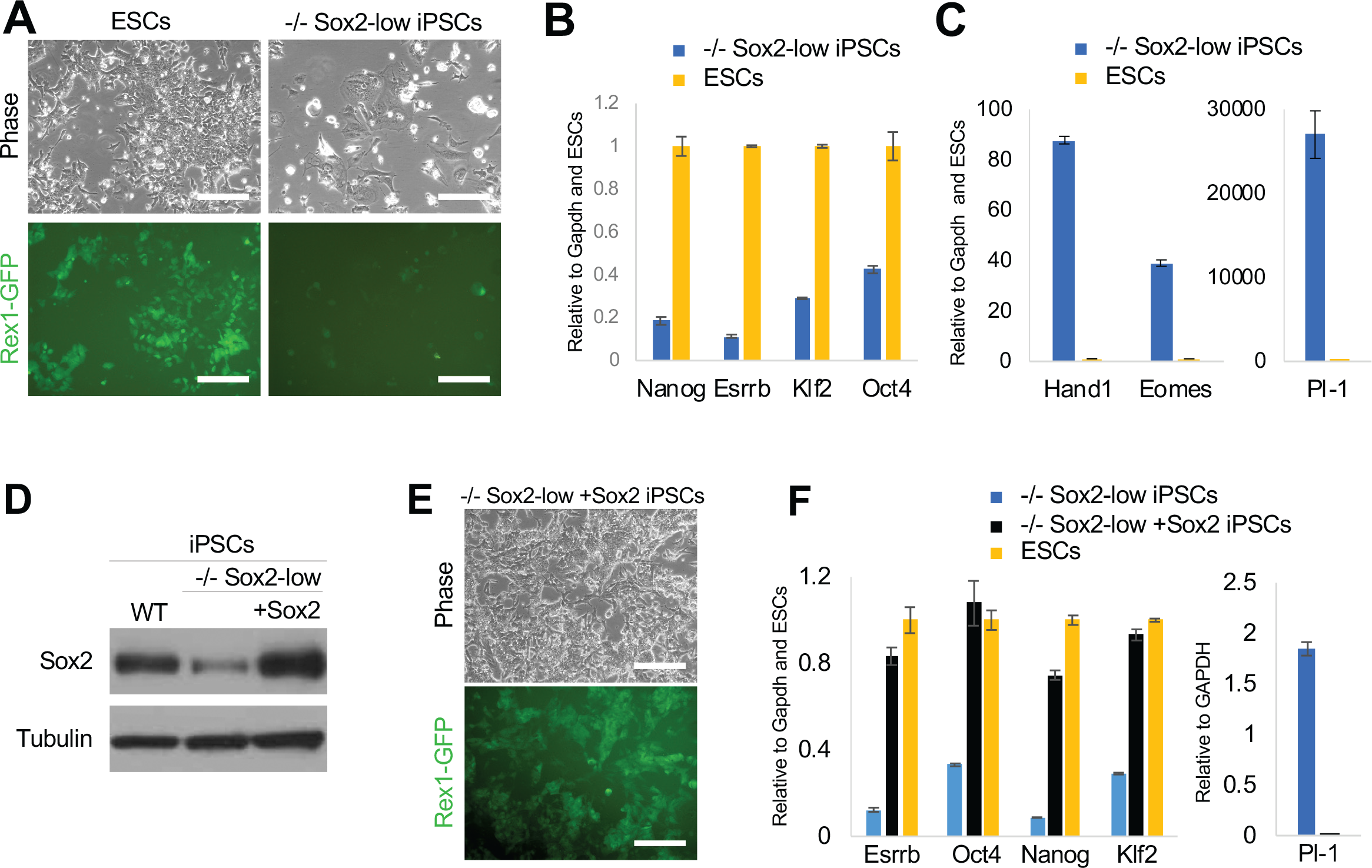
Sox2-low iPSCs do not self-renew in serum plus LIF. A) Phase and Rex1-GFP images of *Sox2−/−* rMKOS (−/− Sox2-low) iPSCs and ESCs at passage 0 after medium switch from 2i plus LIF into serum plus LIF. B-C) qRT-PCR analysis of pluripotency markers (B) and trophoblast markers (C) 3 days after switching into serum plus LIF medium. D) Western blot of Sox2-low iPSCs in 2i plus LIF with and without rescue Sox2 transgene (+Sox2). Rex1-GFP WT iPSCs were used as control for Sox2 protein levels. E) Phase and GFP images of −/− Sox2-low +Sox2 rescue iPSCs after 5 passages in serum plus LIF. F) qRT-PCR analysis, after 3 days in serum plus LIF, of pluripotency factors and Pl-1 in −/− Sox2-low iPSCs with and without a rescue Sox2 transgene. ESCs were provided as control in the qRT-PCR analysis of pluripotency factors. Scale bars = 200µm. Error bars indicate standard deviation of replicate qPCR reactions (n=3).

These results suggest that under weaker self-renewing culture conditions low levels of Sox2 are not sufficient to sustain a naïve pluripotent identity. It also showed that at least a proportion of the cells differentiate towards the trophoblast lineage.

### Reduced Sox2 expression is associated with increased plasticity *in vitro*

To functionally characterize the potency of Sox2-low iPSCs we investigated their differentiation capacity in embryoid body (EB) assays. These efficiently downregulated pluripotency genes and upregulated markers of all 3 germ layers including the ectoderm marker FGF5. Importantly, retroviral Sox2 was not upregulated during differentiation (Figure 3B). Because of the observed upregulation of trophoblast markers in Sox2-low iPSCs in SLIF we also explored the expression of these. In contrast to control lines, Sox2-low iPSCs upregulated expression of trophoblast markers during EB differentiation (Figure 3A). This phenotype was fully rescuable upon the restoration of Sox2 protein levels by the transfection of a constitutive Sox2 transgene (Figure 2D and 3C). These results suggest that Sox2-low iPSCs have increased potency as they can differentiate into both embryonic and extraembryonic lineages.

**Figure 3.**
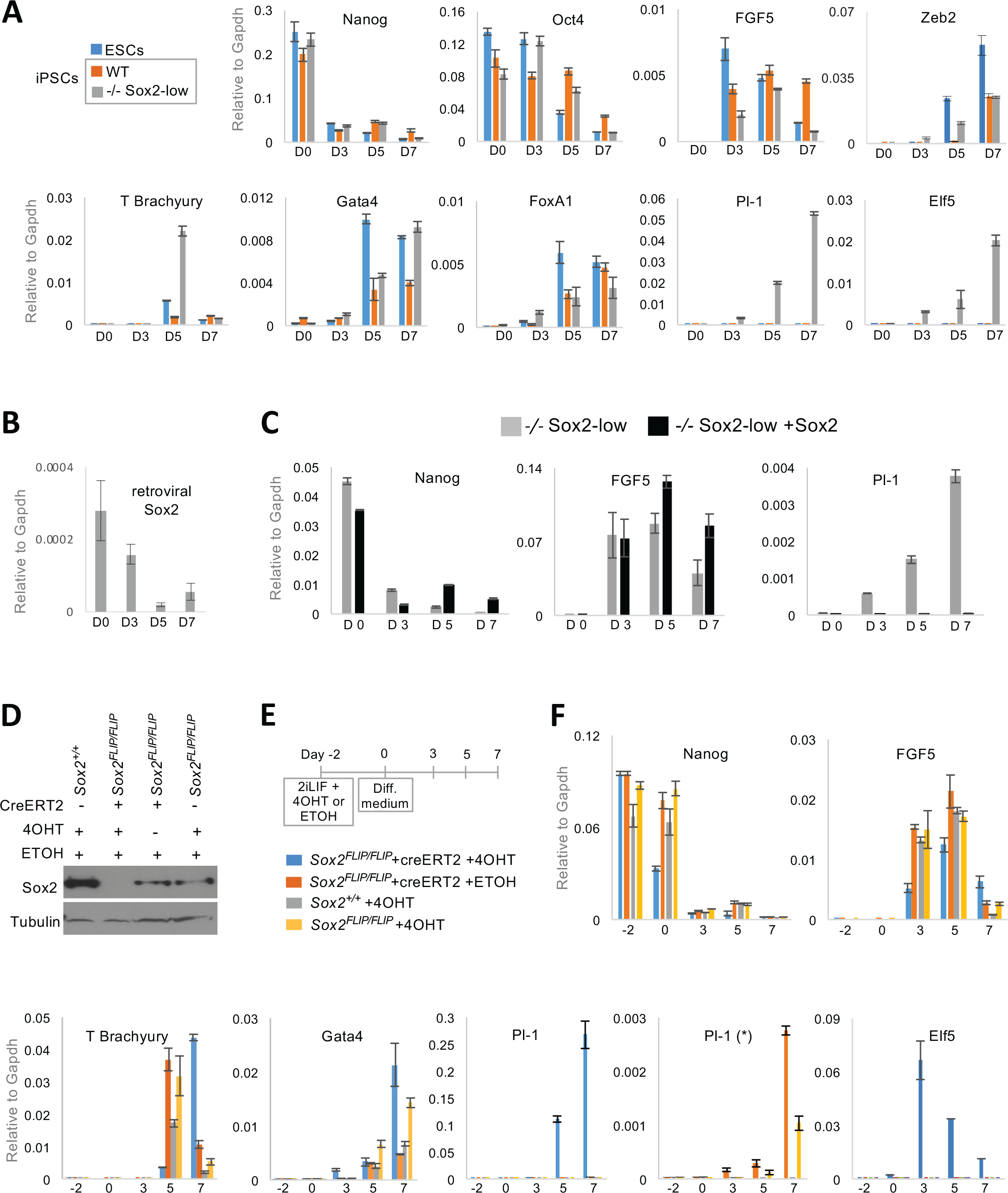
Sox2-low and null nPSCs differentiate into extraembryonic and all embryonic lineages. A) qRT-PCR analysis of embryoid body assay using *Sox2−/−* (−/−) Sox2-low iPSCs, control iPSCs (WT) and ESCs showing RNA expression of markers of pluripotency (Nanog, Oct4), ectoderm (FGF5), mesoderm (T Brachyury and Zeb2), endoderm (FoxA1 and Gatat4) and trophoblast (Pl-1 and Elf5). B) qRT-PCR analysis of embryoid body assay for retroviral Sox2 expression in −/− Sox2-low iPSCs. C) qRT-PCR analysis of embryoid body assays of −/− Sox2-low and rescue (+Sox2) iPSCs, for expression of pluripotency (Nanog), ectoderm (FGF5) and trophectoderm (Pl-1) markers. D) Western blot showing Sox2 protein post tamoxifen (4OHT) treatment in *Sox2FLIP/FLIP* ESCs with or without a constitutive CreERT2 transgene. Carrier only (ethanol, ETOH) and *Sox2+/+* ESCs were used as additional controls. E) Experimental design of embryoid body assay and of Sox2 deletion in *Sox2FLIP/FLIP* ESCs. F) qRT-PCR analysis of embryoid body assay using Sox2FLIP/FLIP ESCs showing expression of pluripotency markers (Nanog), ectoderm (FGF5), mesoderm (T Brachyury), endoderm (Gata4), trophoblast (Pl-1 and Elf5). Pl1 (*) chart omits *Sox2FLIP/FLIP* CreERT2 ESC sample treated with 4OHT. Error bars indicate standard deviation of replicate qPCR reactions (n=3).

Next we went on to test our findings in an independent naïve pluripotent stem cell system (Figure 3D-F and Figure S2A-C). *Sox2FLIP/FLIP* ESCs have previously been generated, which delete endogenous Sox2 expression upon exposure to Cre (Figure S2A) (Andersson-Rolf et al., 2017). Notably, *Sox2FLIP/FLIP* ESCs express Sox2 protein at a reduced level compared to WT ESCs (Figure 3D). This is likely the result of the inserted cassette sequences interfering with the fine regulation of endogenous Sox2. To permit inducible Sox2 deletion, *Sox2FLIP/FLIP* ESCs were constitutively transfected with creERT2. These were then subjected to a tamoxifen timecourse to establish the earliest timepoint at which Sox2 protein is absent. By 36 hours Sox2 protein was undetectable, whereas Nanog and Oct4 were still expressed at high levels at 48 hours, suggesting that differentiation had not occurred (Figure S2B, C). Therefore, to investigate the potency of Sox2-null ESCs we treated CreERT2 *Sox2FLIP/FLIP* ESCs with tamoxifen (4OHT) for 48 hours and then performed EB differentiation (Figure 3D-F). The absence of Sox2 protein did not affect the expected kinetics of downregulation of pluripotency markers or the upregulation of differentiation markers of all 3 germ layers (Figure 3E). Importantly, absence of Sox2 expression was also associated with a striking upregulation of trophoblast markers Pl-1 and Elf5 (Figure 3F). Interestingly, parental *Sox2FLIP/FLIP* ESCs, which exhibit lower Sox2 protein expression compared to control ESCs, also exhibit significant upregulation of the trophoblast marker Pl-1 compared to control ESCs (Figure 3F).

Importantly, these results confirm that reduced Sox2 expression in nPSCs (iPSCs and ESCs) is associated with a gain in cell potency, that is, an ability to differentiate towards the extraembryonic trophoblast lineage in addition to the embryonic lineages.

### Low Sox2 expression does not impair neurectoderm differentiation

Sox2 is thought to be required to drive ectoderm differentiation in pluripotent cells (Thomson et al., 2011). Interestingly, Sox2 null ESCs (Sox2fl/fl pre-treated with Cre) and Sox2-low iPSCs showed upregulation of the ectoderm marker FGF5 in EB assays that was comparable to respective controls (Figure 3A, C and F). To examine the neurectoderm differentiation potential of these cells in more detail, we performed a neural monolayer differentiation (Ying et al., 2003). The Sox2-low iPSCs upregulated neural markers Sox1, Pax6 and Ascl1 to a similar level to WT iPSCs and Sox2-low rescue iPSCs, and gained the characteristic neural rosette morphology, demonstrating efficient neural differentiation (Figure 4A and C). This also occurred without an increase of retroviral Sox2 expression (Figure 4B). These results suggest that reduced Sox2 expression does not impair robust neural differentiation *in vitro*.

**Figure 4.**
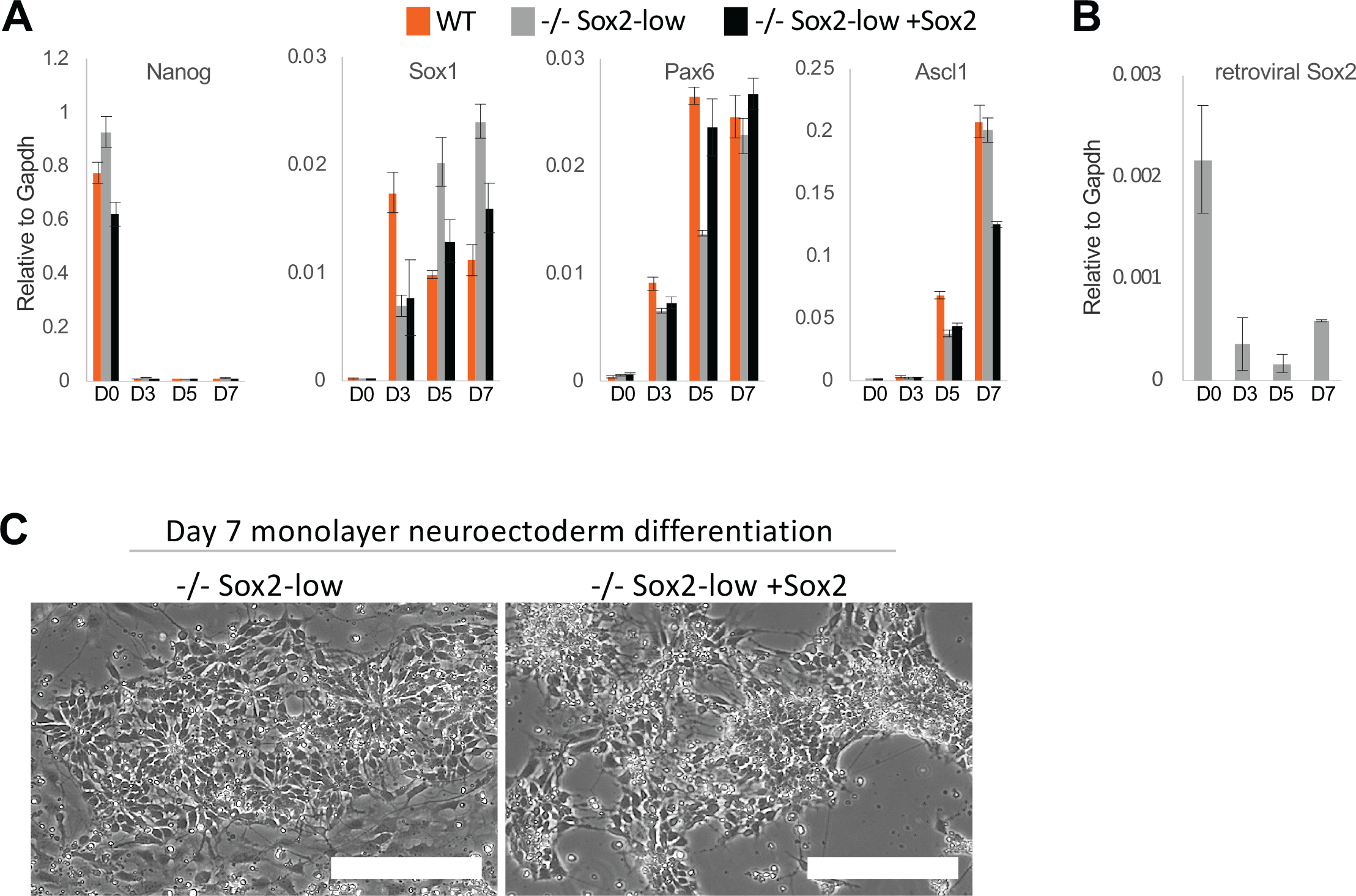
Low Sox2 expression is compatible with robust neuroectoderm differentiation. A) qRT-PCR analysis of *Sox2−/−* (−/−) Sox2-low, −/− Sox2-low rescue (+Sox2) and WT iPSCs in a neurectoderm monolayer differentiation assay for markers of pluripotency (Nanog) and neurectoderm (Pax6, Ascl1, and Sox1). B) qRT-PCR analysis of retroviral Sox2 in −/− Sox2-low iPSCs during neurectoderm monolayer differentiation. C) Phase images of −/− Sox2-low iPSCs and Sox2-low rescue iPSCs on day 7 of the neurectoderm monolayer differentiation assay. Error bars indicate standard deviation of replicate qPCR reactions (n=3).

### Reduced Sox2 expression is associated with increased plasticity *in vivo*

To investigate the effect of reduced Sox2 expression in differentiation *in vivo*, we performed morula aggregations with Sox2-low iPSCs. To visualize embryo chimaerism we constitutively transfected Sox2-low iPSCs with MST-dsRed fluorescent protein. Consistent with the findings *in vitro*, there was contribution of Sox2-low iPS-derived cells to both epiblast and presumptive trophoblast at E7.5 (Figure 5A). In order to determine trophoblast chimaerism we also generated Sox2-low and control iPSCs expressing constitutive GFP. As expected, E6.5 WT iPSCs contributed exclusively to the epiblast (Figure 5B-D). In contrast Sox2-low iPS-derived cells were found in both the embryonic and extraembryonic, as defined by the AP2γ protein domain, compartments of the embryos (Figure 5B, C and E and S3A). Lineage marker staining showed that 83% of Sox2-low chimaeras exhibit contribution to both the trophoblast and epiblast compartments (Figure 5E). In fact 90% of all Sox2-low chimeras showed trophoblast contribution. Interestingly, some Sox2-low iPS-derived cells exhibited co-expression of trophoblast marker AP-2γ and epiblast marker Oct4 (filled arrowheads, Figure 5C) and this occurred in both the epiblast and trophoblast embryo compartments.

**Figure 5.**
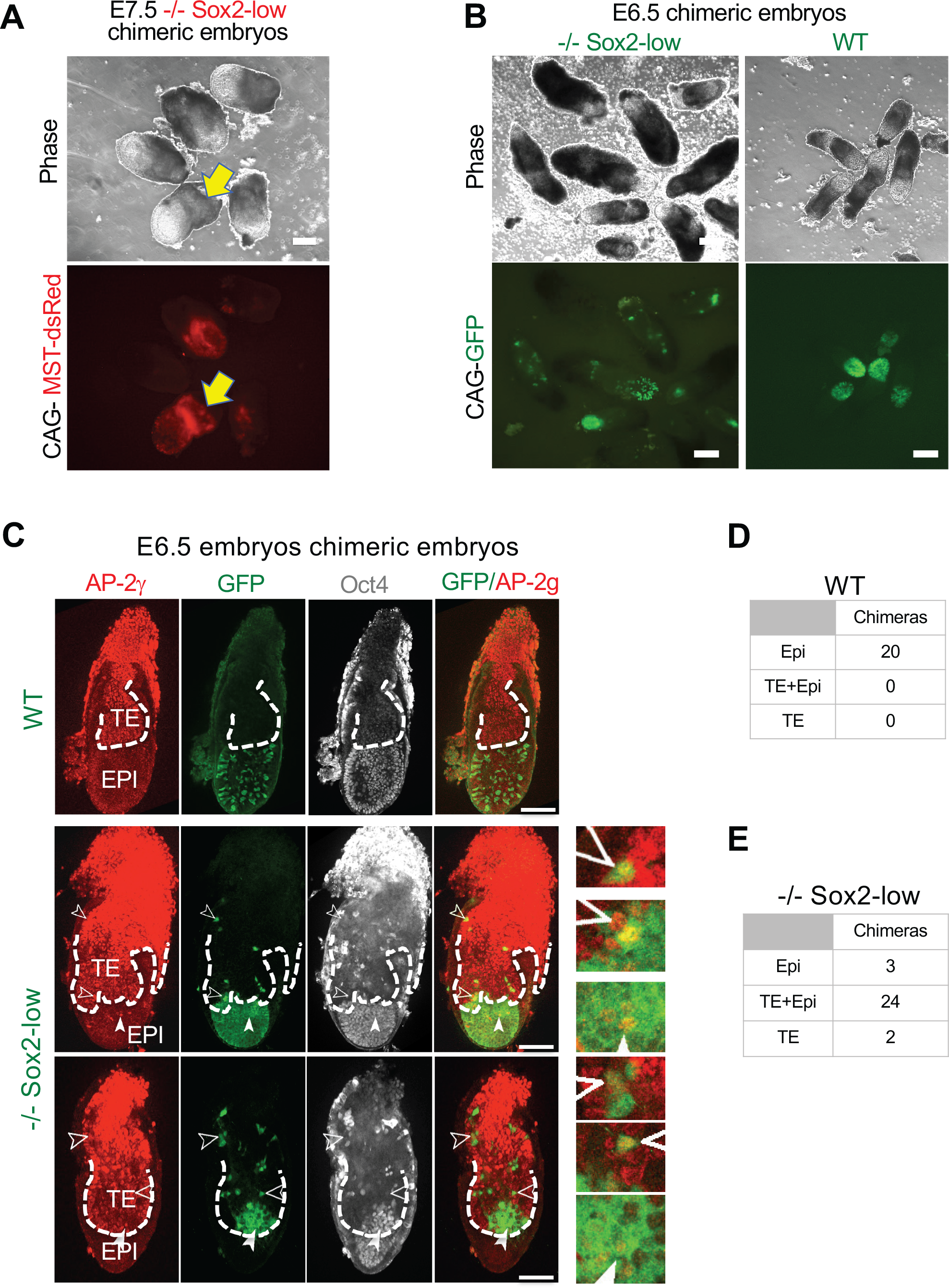
Sox2-low iPSCs exhibit expanded potency in vivo. A) Phase and red fluorescence images of E7.5 chimaeras of *Sox2−/−* (−/−) Sox2-low iPSCs expressing constitutively MST-dsRed (red fluorescence). Arrow indicates presumptive contribution to the trophoblast/trophectoderm lineage. Scale bars = 200µm. B) Phase and GFP images of E6.5 chimeric embryos generated with either −/− Sox2-low or *WT* iPSCs constitutively expressing a GFP transgene. Please note that apparent difference in size is due to −/−Sox2 chimeric embryos having been imaged with some maternal tissue still attached. Scale bars = 200µm. C) Single confocal microscopy sections of indicated genotype chimeric E6.5 embryos stained with trophoblast (AP-2γ) and epiblast (Oct4) markers. Filled arrowheads indicate examples of chimeric cells co-expressing Oct4 and AP-2γ markers. Non-filled arrowheads indicate examples of chimeric cells expressing AP-2γ only. Epiblast (EPI) and Trophoblast/trophectoderm (TE) embryo domains are separated by dashed line. Scale bars = 100µm. D-E) Table showing compartmental contribution of *WT* (D) or −/− Sox2-low (E) iPSCs constitutively expressing a GFP transgene.

To define when Sox2-low iPS derived cells start contributing to the trophoblast lineage we injected these at the morula stage and assessed chimaerism at the blastocyst stage. Interestingly, almost all of the chimaeras showed contribution to the epiblast only (Figure S3B and C). This data supports *in vitro* assays showing that Sox2-low iPSCs only acquire trophoblast lineage differentiation potential when in an environment no longer supportive of naïve pluripotent cell identity.

Overall, these data demonstrate that Sox2-low iPSCs are competent to contribute to both embryonic and extraembryonic embryo development.

### Sox2-low iPSCs have a basal TE molecular signature

To ascertain the identity of Sox2-low iPSCs we performed single-cell RNAseq on these and control WT and Sox2-low +Sox2 rescue iPSCs. Gene ontology enrichment for genes differentially expressed between Sox2-low iPSCs and control iPSCs revealed only 325 differentially expressed genes between the different genotypes with no strongly enriched GO terms (Figure S4). In addition, no Sox family member gene was found upregulated in Sox2-low iPSCs, eliminating that way possible functional redundancy (Corsinotti et al., 2017). In agreement with Sox2-low iPSCs having a naïve identity they clustered closely with control iPSCs, ESCs and E4.5 naïve epiblast cells and separate from embryo trophoblast/trophectoderm (TE) cells (Figure 6A). However, when looking at the identity of single iPSCs on a continuum from an embryonic TE or naïve epiblast signature Sox2-low cells were generally found in the fractions with lower epiblast and higher TE molecular identities (Figure 6B). We also looked at accumulative gene expression for genes known to be associated with either the TE or with the inner cell mass (ICM) of embryos (Blakeley et al., 2015) (Figure 6C). Again, this revealed that Sox2-low cells can be distinguished from WT and rescue lines as they display higher TE and lower ICM gene expression.

**Figure 6.**
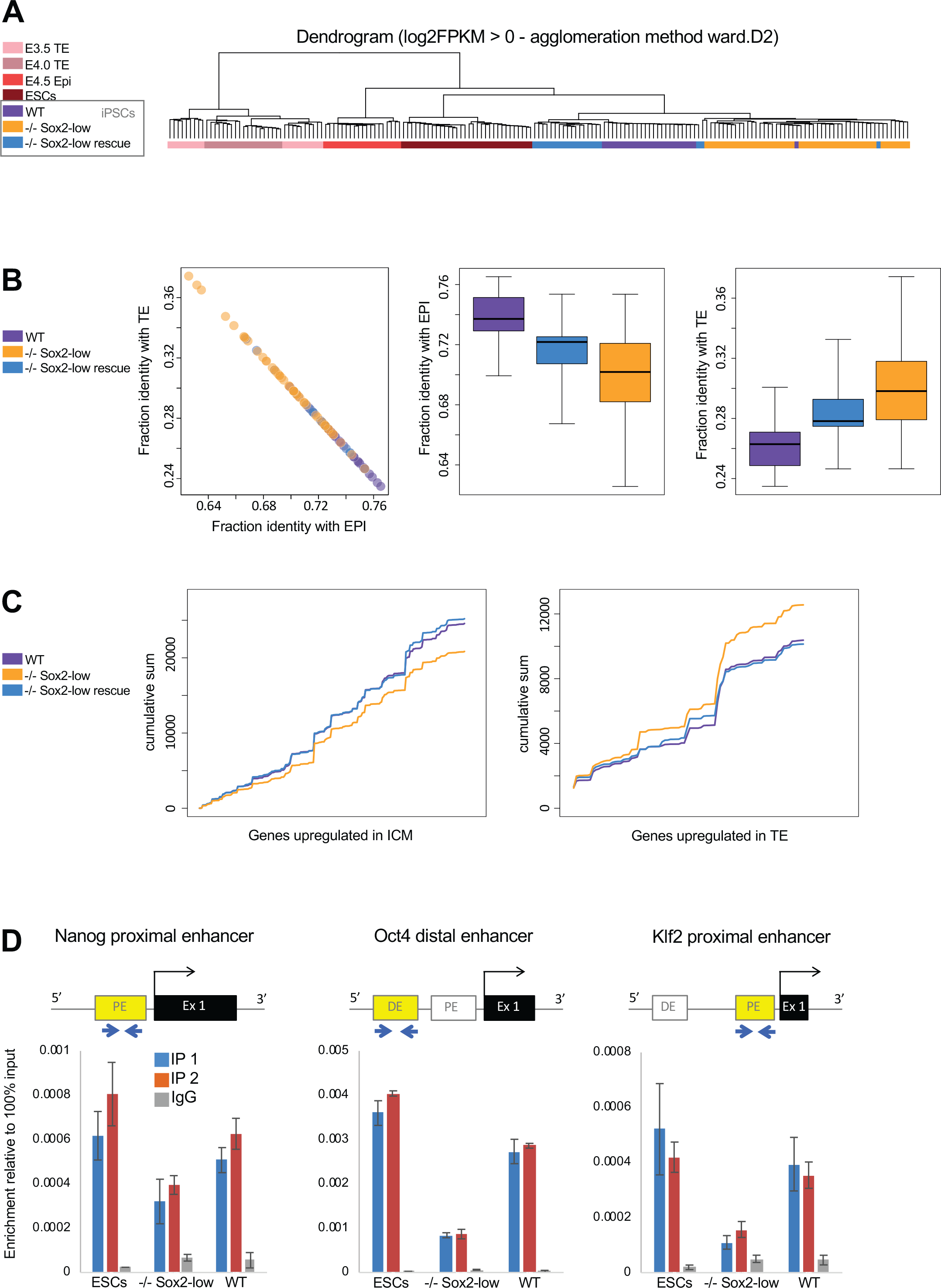
Sox2-low iPSCs display a basal TE molecular signature. A) Dendrogram computed with the top variable genes among the selected cell types (FPKM > 1, logCV2 > 0.5, n=1446). Trophectoderm (TE) E3.5 and E4.0 Single-cell RNA sequencing (scRNA-seq) data is from (Deng et al., 2014), E4.5 Epiblast (Epi) scRNA-seq data is from (Mohammed et al., 2017), scRNA-seq ESC data is from (Sousa et al., 2018). B) Scatterplot of fraction of identity between WT, −/− Sox2-low, −/− Sox2-low rescue iPSC line samples and embryo lineages (epiblast and trophectoderm). Boxplot of the distribution of the fraction of identity between each population. C) Cumulative sum for genes upregulated in TE or ICM (Blakeley et al., 2015), computed with gene expression value for WT, −/− Sox2-low, −/− Sox2-low rescue iPSC line samples. D) Oct4 chromatin immunoprecipitation in Rex1-GFP ESCs and iPSCs in 2iLIF. IP = immunoprecipitation. IgG = negative control. Error bars represent the standard deviation of three technical qPCR replicates. IP1 and IP2 represent independent Oct4 immunoprecipitations. IgG represents negative control normal IgG immunoprecipitation.

These results suggest that observed increased potency of Sox2-low cells is a functional property present at the single cell level, as opposed to cell population heterogeneity. This may also be linked to the underlying basal TE molecular signature.

### Reduced Sox2 expression impairs Oct4 DNA-binding

Sox2 is thought to be a DNA-binding partner to Oct4, and Oct4 inhibits extraembryonic differentiation (Niwa et al., 2000; Reményi et al., 2003). Therefore, we hypothesised that reduced Sox2 protein level may impact directly or indirectly on Oct4 DNA-binding ability in naïve pluripotent stem cells. Consistent with this notion, Oct4 chromatin immunoprecipitation showed reduced Oct4 genomic occupancy in Sox2-low iPSCs at key naïve pluripotent regulatory sequences (Figure 6D).

These results suggest that reduced Oct4 occupancy at naïve associated regulatory sequences combined with the basal TE molecular signature are likely underlying causes rendering Sox2-low iPSCs competent to also contribute to the trophoblast lineage.

## Discussion

Here we show that reduced levels of Sox2 are compatible with self-renewal of nPSCs in 2iL conditions, although they collapse in SLIF demonstrating fragility in their pluripotent network. This fragility is likely due to the basal TE molecular identity and to the reduced Oct4 DNA-binding at naïve-associated regulatory sequences.

Sox2 and Oct4 have been hypothesised to bind together at Oct/Sox elements to drive the pluripotency network (Chew et al., 2005; Rodda et al., 2005). It has been shown that Sox2 binds to the DNA target sequence first and that it recruits Oct4 (Chen et al., 2014). However, Oct4 and Sox2 were also reported to operate in a largely independent manner and their impact on each-others ability to bind to regulatory sequences may be indirect and regulated by chromatin accessibility (Friman et al., 2019). Independent of the Oct4-Sox2 relationship these studies are consistent with our findings that reduced Sox2 expression disrupts optimal Oct4 DNA-binding.

We found that reduced or complete depletion of Sox2 is associated with differentiation into extraembryonic lineages, as well as the 3 germ layers. This indicates that Sox2 plays a role in reducing the potency of nPSCs such that they cannot differentiate into extraembryonic lineages. The mechanism by which reduced Sox2 expression levels releases an inhibition on extraembryonic differentiation is likely linked to the impaired Oct4 genomic occupancy. Similar to Sox2 deletion, Oct4 deletion in ESCs causes differentiation into trophectoderm-like cells (Niwa et al., 2000). However, unlike Sox2-low nPSCs, which can contribute to both embryonic and extraembryonic tissues, Oct4-low nPSCs lose the ability to differentiate into embryonic lineages (Radzisheuskaya et al., 2013). This suggests that in contrast to Oct4, Sox2 is not a key factor regulating differentiation towards embryonic lineages.

The importance of Sox2 in inhibiting extraembryonic differentiation is counter to the evidence suggesting Sox2 has a key role in extraembryonic development. Sox2 is required to derive trophoblast stem cells and is expressed in extraembryonic tissues post-implantation (Avilion et al., 2003). However, this is likely to be an *in vitro* only requirement as *Sox2−/−* extraembryonic tissues are sufficient for development at least up to E12.5 (Avilion et al., 2003). Morula aggregations using the Sox2-low iPSCs showed that only one out of 28 chimeric embryos exhibited trophectoderm contribution by the blastocyst stage. This is in stark contrast to E6.5 at which point 90% of all chimeric embryos display trophectoderm contribution. Interestingly, we observed cells within the epiblast upregulating the extraembryonic marker AP-2γ. This is in agreement with Sox2-low iPSCs being naive, but upon differentiation showing additional competency to upregulate TE lineage markers and to contribute towards this lineage. Thus, their ability to undergo an extraembryonic lineage fate occurs after naïve pluripotent cell identity exit and simultaneously with the embryonic lineage fate. The already reduced Oct4 genomic occupancy combined with the onset of the downregulation of naïve factors upon initiation of cell differentiation may somehow create a window of opportunity for differentiating naïve cells to also acquire a TE fate.

Recently, expanded potential stem cells (EPSCs) that can contribute to both embryonic and extraembryonic lineages have been generated by two independent laboratories, by culturing ESCs in the presence of small molecules (Yang et al., 2017a, 2017b). The two laboratories used different chemical cocktails to generate the EPSCs but both resulted in cells with similar properties, suggesting a possible common mechanism. Among the added chemicals are minocycline hydrochloride (Yang et al., 2017b), a Parp1 inhibitor, and XAV939 (Yang et al., 2017a), a tankyrase inhibitor. The contributory mechanism of XAV939 to the extension of pluripotency is unclear, but it likely inhibits Parp family members TNKS1/2 and/or stabilises AXIN (Yang et al., 2017a). Furthermore, Parp1−/− ESCs have a propensity to differentiate into trophoblast (Hemberger et al., 2003). Parp1 is thought to aid Sox2 binding in ESCs (Liu and Kraus, 2017). Therefore, the previously published EPSC culture condition may reduce PARP activity and consequently reduce Sox2 DNA-binding, thus resulting in a similar phenotype to our Sox2-low nPSCs.

In conclusion our study identifies an unexpected role of a naïve pluripotency factor as a restrictor of potency and provides a conciliatory mechanistic explanation for EPSCs, which exhibit both embryonic and extraembryonic potency.

## Author contributions

J.C.R.S. conceived and supervised the study, designed experiments, wrote and approved the manuscript. K.T. designed and performed experiments, analyzed the data and wrote the manuscript. A.R. initiated the study, designed experiments and helped in the supervision. L.E.B. designed and performed the Oct4 ChIP experiment. K.M. performed the neuroectoderm monolayer and other experiments. K.J. performed embryo dissections. A.A.R. generated *Sox2FLIP/FLIP* ESCs overseen by B.K.K. H.T.S. designed and performed the single cell sequencing experiment. G.G.S. and P.B. designed and performed bioinformatic analyses.

## Acknowledgements

We thank Yael Costa for critical reading of the manuscript. Peter Humphreys for assistance with imaging. William Mansfield for blastocyst injections. This study was supported by a Wellcome Trust Fellowship (WT101861) to J.C.R.S., who is a Wellcome Trust Senior Research Fellow. B.K.K is supported by a European Research Council grant (639050). K.M. is a recipient of a Darwin Trust of Edinburgh Ph.D. studentship. K.T. is a recipient of a MRC Ph.D. studentship. H.T.S. and L.E.B. were supported by MRC research grant MR/R017735/1. G.G.S. is funded by BBSRC research grant RG53615.

---

**Figure S1.**
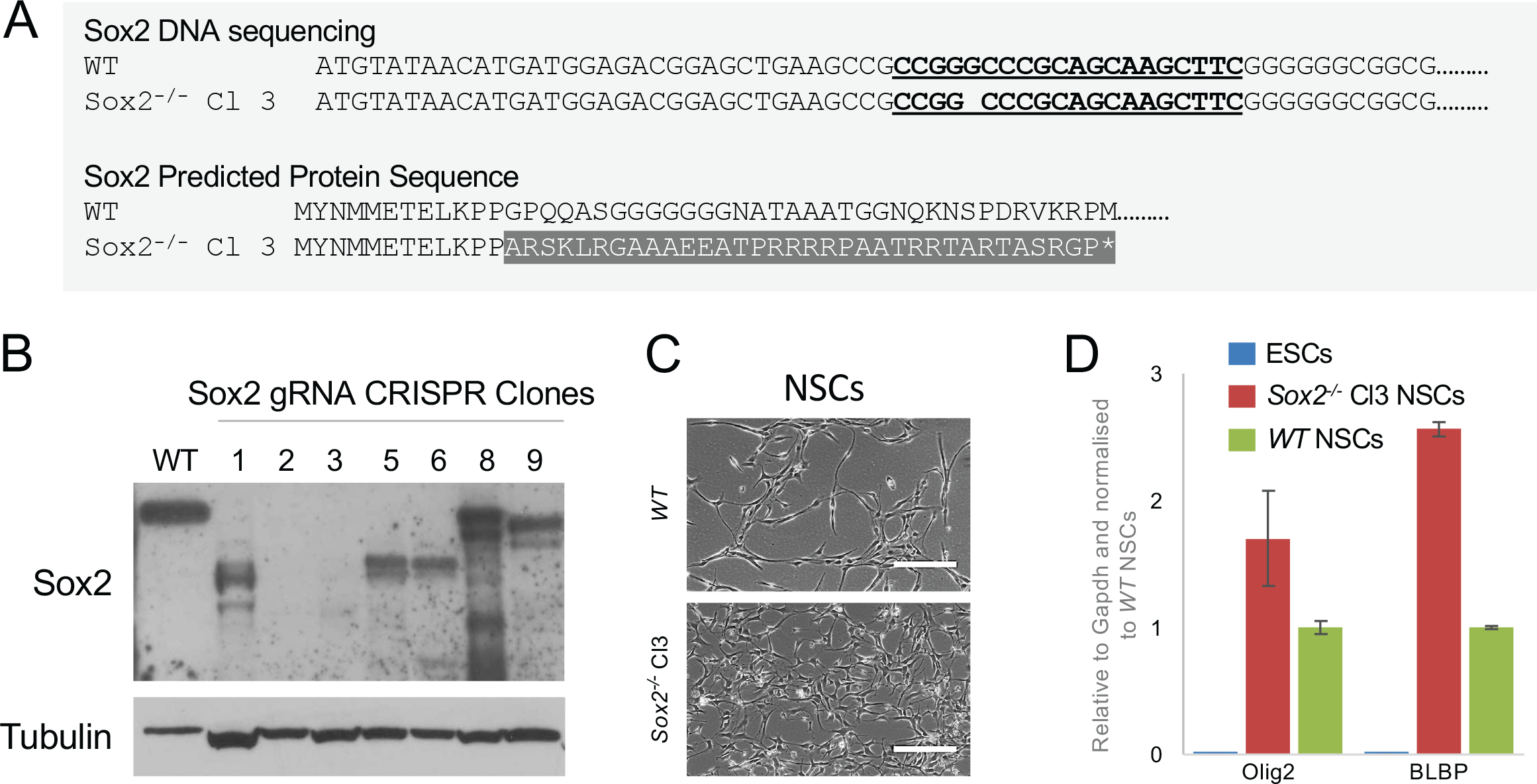
Generation of *Sox2−/−* NSCs. Related to Figure 1. A) Sequence of start of coding region of *Sox2* amplified by genomic PCR for *Sox2−/−* neural stem cell (NSC) clone 3 with Cas9 targeting gRNA binding site underlined aligned with the WT Sox2 coding region to show the mutation (deletion of 1 guanine). The predicted protein sequence from the *Sox2−/−* NSC clone 3 is shown aligned with WT sequence, with the sequence diversion highlighted. * = stop codon. B) Western blot for Sox2 (≈40kDa) and Tubulin (≈50kDa) protein expression in WT and clonal NSC lines after transfection with Sox2 gRNA/Cas9. C) Phase images of *WT* and *Sox2−/−* clone (Cl) 3 NSCs. D) qRT-PCR analysis of neural markers (Olig2 and BLBP) in *WT* and *Sox2−/−* Cl3 NSCs and ESCs. Error bars indicate standard deviation of replicate qPCR reactions (n=3). Scale bars = 200µm.

**Figure S2.**
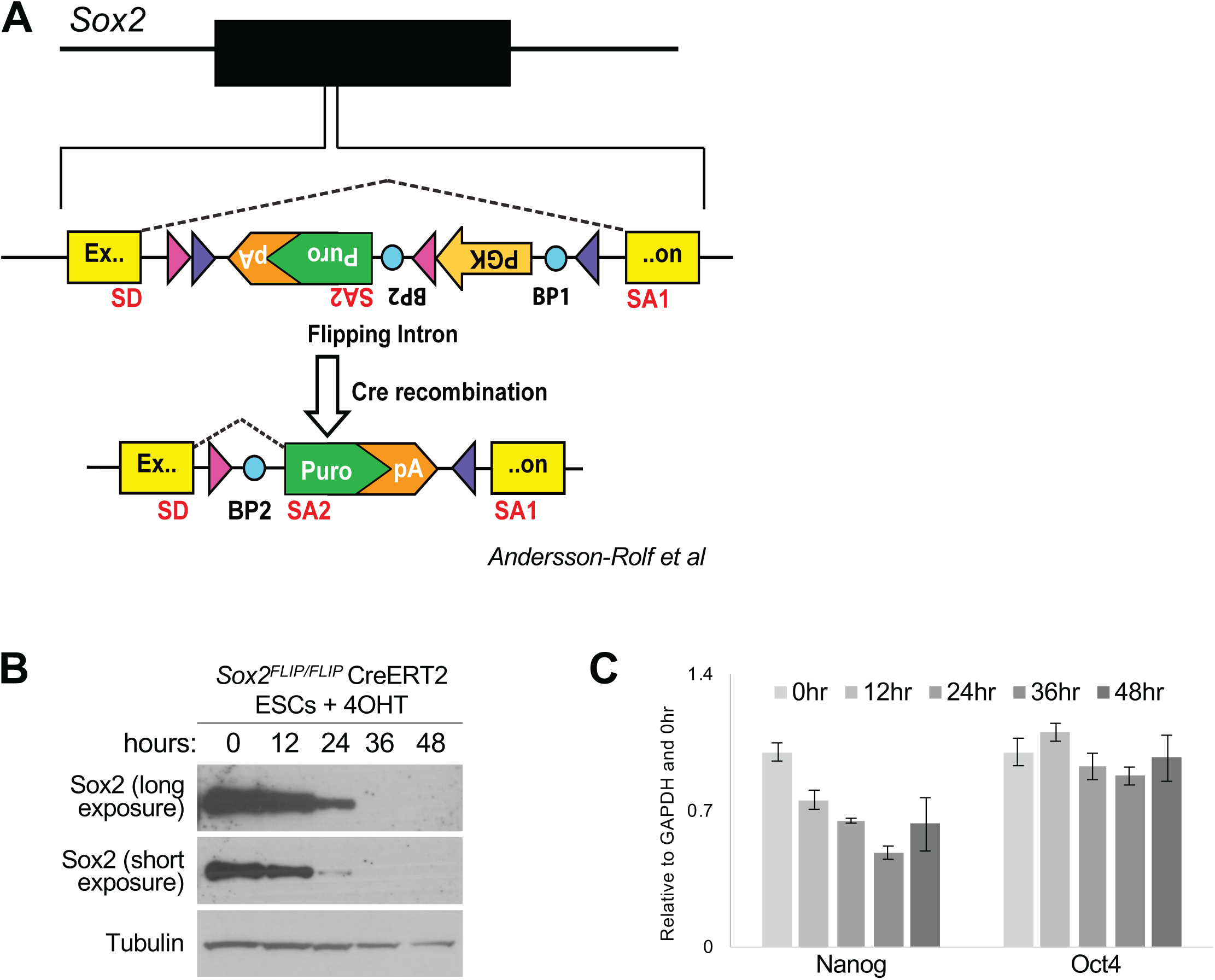
*Sox2FLIP/FLIP* ESCs experimental optimisation. Related to Figure 3. A) Schematic of Sox2FLIP/FLIP ESCs. Please note that after Cre mediated recombination Sox2 protein becomes truncated. SD indicates splice donor site. BP1 and SA1, and BP2 and SA2 indicate pairs of branch points and splice acceptor sites that are used before and after Cre recombination respectively. Pink and purple triangles indicate LoxP variants (Lox5171 and LoxP1 respectively). Dashed line indicates splice site usage. B) Western blot of 48 hour timecourse of tamoxifen (4OHT) treatment of Sox2FLIP/FLIP CreERT2 ESCs. C) qRT-PCR of pluripotency factor expression during tamoxifen timecourse. Error bars indicate standard deviation of replicate qPCR reactions (n=3).

**Figure S3.**
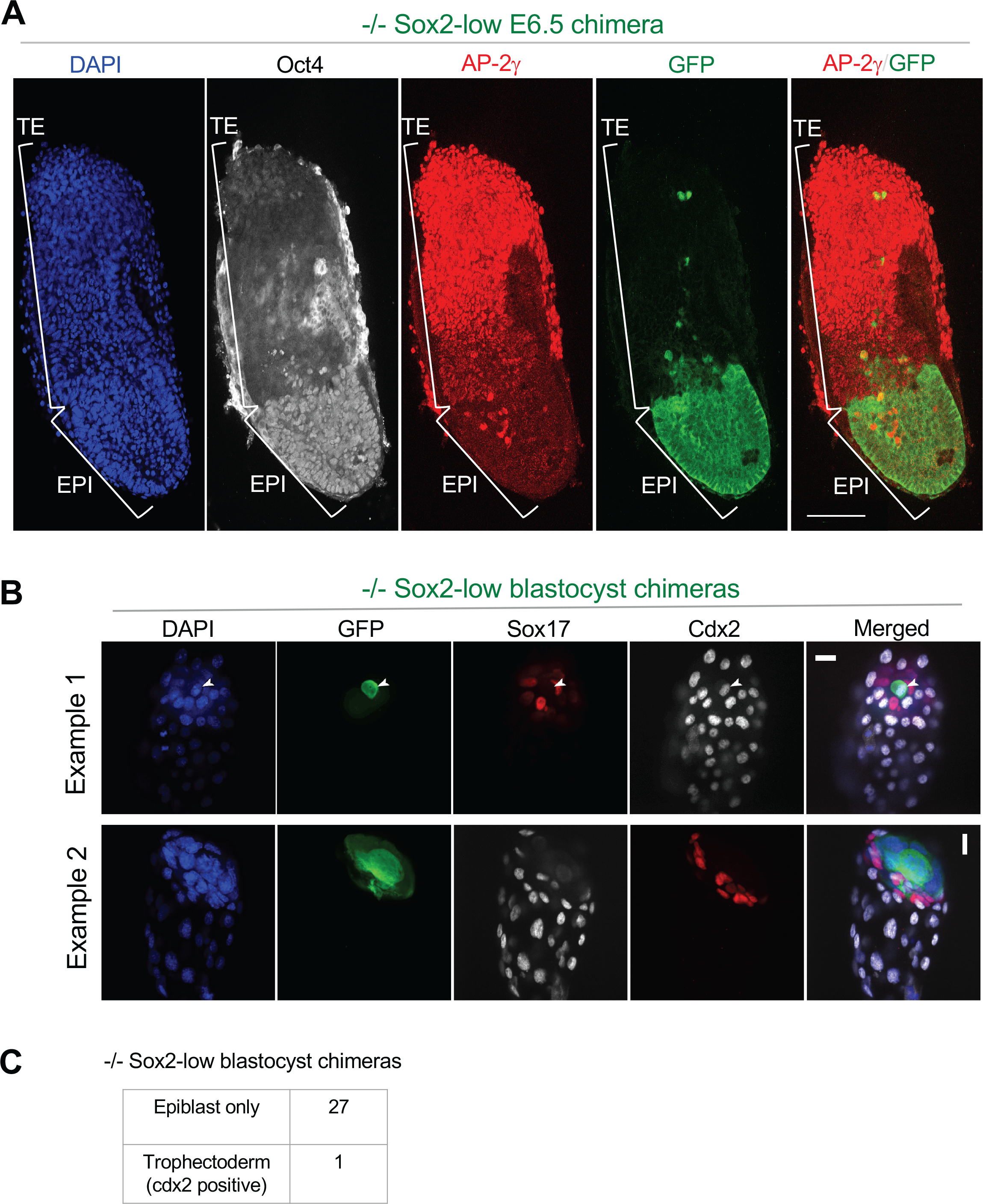
Sox2-low iPSCs exhibit expanded potency in vivo. Related to Figure 5. A) Single confocal microscopy section of *Sox2−/−* (−/−) Sox2-low iPSCs (GFP) E6.5 chimeric embryos stained with trophoblast (AP-2**γ**) and epiblast (Oct4) markers. Epiblast (EPI) and Trophoblast/trophectoderm (TE) embryo domains are indicated. Scale bars = 100**μ**m. B) Immunofluorescence staining with Cdx2 (trophectoderm) and Sox17 (hypoblast) of cultured embryos after morula injection with *Sox2−/−* (−/−) Sox2-low iPSCs expressing constitutively a GFP transgene. Scale bar = 20**μ**m. C) Table showing compartmental contribution of −/− Sox2-low iPSCs.

**Figure S4.**
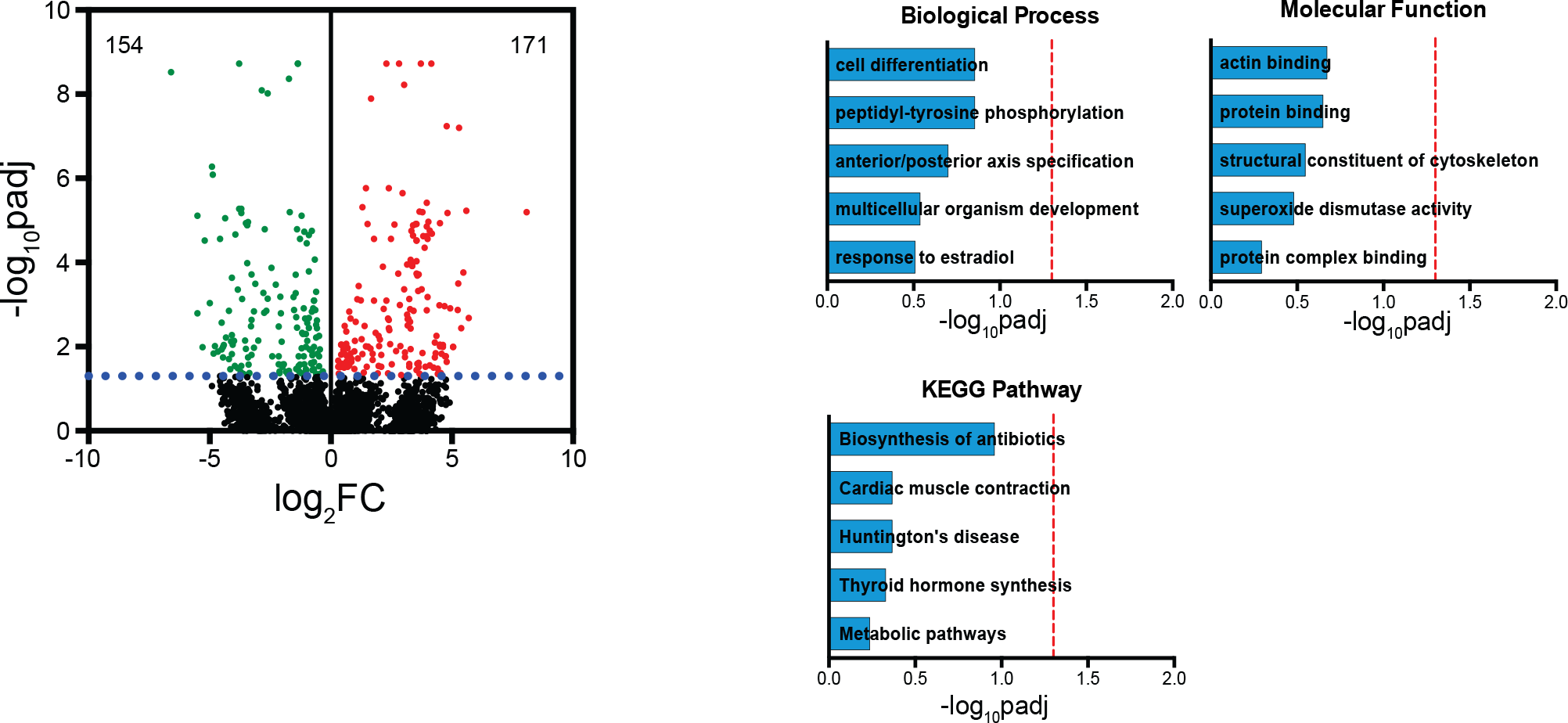
Sox2-low iPSCs do not display gene ontology differences to control cell lines. Volcano plot and gene ontology enrichment for genes differentially expressed between *Sox2−/−* (−/−) Sox2-low and control iPSCs, which include WT and −/− Sox2-low rescued with constitutive Sox2 transgene (+Sox2). Significance threshold was defined at ‑log_10_padj=1.3.

## Material and methods

### Plasmids

pMXs c-Myc, pMXs Oct4, pMXs Sox2, pMXs Klf4 and pSpCas9(BB)-2A-GFP (Addgene). pPyCAG-MST-IRES-Puro, pPyCAG-eGFP-IRES-Zeo (Austin Smith). pPB-CAG-DEST-PGK-hph, pCAG-CreERT2NLS-IRES-bsd (Joerg Betschinger).

### Cell culture

PLAT-E cells and (where stated) iPSCs were cultured in GMEM basal medium (GMEM (Sigma-Aldrich), 1xNEAA (Gibco), 1xpenicillin-streptomycin (Sigma-Aldrich), 1mM sodium pyruvate (Sigma-Aldrich), 0.1mM 2-mercaptoethanol (ThermoFisher Scientific) and 2mM L-glutamine (ThermoFisher Scientific)) with 10% FCS (Labtech) and 20ng/ml mouse LIF (University of Cambridge). NSCs were cultured in DMEM/F12 (ThermoFisher Scientific), 1xNEAA, 0.1mM 2-mercaptoethanol, 1xpenicillin-streptomycin, 1:100 v/v B27 supplement (ThermoFisher Scientific), 1:200 v/v N2 supplement (University of Cambridge), 4.5μM HEPES (ThermoFisher Scientific), 0.03M glucose and 120μg/ml BSA (ThermoFisher Scientific) supplemented with 10ng/ml EGF (Peprotech) and 20ng/ml FGF2 (University of Cambridge). iPSCs were cultured in KSR basal medium (GMEM basal medium with 10% KSR, 1% FCS) or N2B27 basal medium (DMEM/F12 and Neurobasal (ThermoFisher Scientific) in a 1:1 ratio, 1xpenicillin-streptomycin, 0.1mM 2-mercaptoethanol, 2mM L-glutamine, 1:200 v/v N2 and 1:100 v/v B27 supplement) with additional 20ng/ml mouse LIF, CHIR99021 3µM and PD0325901 1µM (both Biotechnology Center TU Dresden, Stewart lab). Additional chemicals used: hygromycin B 200 µg/ml (ThermoFisher Scientific), blasticidin 40 µg/ml (ThermoFisher Scientific), puromycin 1µg/ml (ThermoFisher Scientific), zeocin 100µg/ml (ThermoFisher Scientific), 4-hydroxytamoxifen 500 nM.

### Reprogramming

Separate PLAT-E cultures were transfected with 9µg of each pMX plasmid using FuGENE 6 (Roche) for retroviral production. 48 hours later the supernatants were collected, combined, and mixed with 4μg/ml Polybrene (Sigma-Aldrich) and filtered through a 0.45 μm cellulose acetate filter. This was applied to NS cells for 24 hours, after which they were returned to NS medium for 2 days, before being placed in serum/LIF to form reprogramming intermediates (preiPSCs). These were then placed in KSR2iLIF to form iPSCs.

### Cell differentiation

For embryoid body assays, 1.5×106 cells were plated in suspension into serum basal media without LIF in 90mm low-attachment dishes for 7 days with media changed every other day.

### Neural differentiation

1×105 cells were plated directly into 6-well plates, previously coated with laminin for 2 hrs at 37 degrees, containing N2B27 plus 1μM Alk inhibitor (A83-01) differentiation medium and then cultured in a low oxygen incubator. Medium was changed every day.

### Western blotting

The primary antibodies used were: rat monoclonal against Sox2 (1:2000, eBioscience, 149811); mouse monoclonal against α-tubulin (1:10000, Abcam, ab7291). The secondary antibodies were HRP-linked antibodies against rat or mouse IgG (GE Healthcare).

### Embryo staining

Embryos were fixed with 4% paraformaldehyde and mounted in vectashield antifade mounting medium (Vector Laboratories). Blastocysts were permeabilised using 100% methanol and blocked using 2% donkey serum, 0.1% BSA and 0.2% Triton X-100. Post-implantation embryos were permeabilised in 0.25% Triton X-100/PBS and blocked using 0.01% Triton X-100/PBS and 3% donkey serum. Primary antibodies used were goat polyclonal against Sox17 (1:500, R and D systems, AF1924), mouse monoclonal against Cdx2 (1:500, Biogenex, MU392A-UC), AP-2γ (1:100, Cell Signalling Technology, 2320), rat monoclonal against GFP (1:200, Nacalai, 04404-26), mouse monoclonal against Oct4 (1:100, Santa Cruz, sc-5279). Secondary antibodies were Alexa Fluor antibodies (ThermoFisher Scientific) and were used at a concentration of 1:500.

### Morula aggregation

E2.5 CD1 embryos were combined with iPSCs and transferred to recipient mice to assess the contribution in post-implantation development or were cultured *in vitro* for 2 days to assess blastocyst contribution.

### Cell transfection

Nucleofection (NSCs) was performed using the AMAXA Nucleofection Technology (Lonza, VCA-1003). Lipofection (ESCs, iPSCs) was performed using Lipofectamine 2000 Transfection Reagent (ThermoFisher Scientific).

### Western Blot Quantification

Quantification was performed using the ‘Gels’ tool in ImageJ Fiji. Background-subtracted relative intensity (BSRI) was calculated for each band of interest. Protein of interest (Sox2) BSRI values were then normalized to the corresponding loading control (tubulin) BSRI values and ESCs.

### Generation of CRISPR/Cas9 cell lines

*Sox2−/−* NSCs were generated by nucleofecting NSCs with 4μg CRISPR/gRNA plasmid and single-cell sorting 48 hours later to generate clonal cell lines. The generation of Sox2 FLIP/FLIP ESCs has been previously described (Andersson-Rolf et al., 2017).

### Chromatin Immunoprecipitation (ChIP)

ChIP was performed as previously described (Radzisheuskaya et al., 2013) with minor changes. Briefly, 1×107 cells were fixed for 10 min in 0.4% formaldehyde then washed with ice-cold PBS. Cells were incubated in lysis buffer 1 (50 mM HEPES pH 7.5, 140 mM NaCl, 1 mM EDTA, 10% Glycerol, 0.5% NP40, 0.25% Tx100) then in lysis buffer 2 (10 mM Tris pH 8.0, 200 mM NaCl, 1 mM EDTA, 0.5 mM EGTA) for 10 min each. Nuclei were resuspended in shearing buffer (1% SDS, 10 mM EDTA, 50 mM Tris pH 8.0) and sonicated to an average fragment size of 300-500 bp. Chromatin was diluted 1:10 in dilution buffer (50 mM Tris pH 8.0, 167 mM NaCl, 1.1% Tx100, 0.11% Na-deoxycholate) and cleared with isotype IgG coated protein G Dynabeads (ThermoFisher Scientific). A portion of chromatin was taken as input control. Chromatin was then incubated overnight at 4°C with 1.5μg Rabbit anti-Oct4 antibody (Abcam; ab19857) antibody or 2μg Rabbit normal IgG (Santa Cruz Biotechnology; sc-2027). Chromatin-antibody mix was incubated with pre-blocked protein G dynabeads for 1 hour at 4°C. These were then washed twice in low salt wash buffer (50 mM Tris pH 8.0, 0.1% SDS, 0.1% Na-deoxycholate, 1% Tx100, 150 mM NaCl, 1 mM EDTA, 0.5 mM EGTA), once in high salt wash buffer (50 mM Tris pH 8.0, 0.1% SDS, 0.1% Na-deoxycholate, 1% Tx100, 500 mM NaCl, 1 mM EDTA, 0.5 mM EGTA), once in LiCl wash buffer (50 mM Tris pH 8.0, 250 mM LiCl, 0.5% Na-deoxycholate, 0.5% NP40, 1 mM EDTA, 0.5 mM EGTA) and twice in TE wash buffer (50 mM Tris pH 8.0, 10 mM EDTA, 5 mM EGTA). Bound chromatin was then eluted at 65°C in elution buffer (1% SDS, 0.1 M NaHCO3). Crosslinking was reversed through overnight incubation at 65°C for samples and inputs. DNA was purified using the QIAquick PCR Purification Kit (Qiagen) according to the manufacturer’s protocol. DNA was quantitated by SYBR Green (ThermoFisher Scientific) qPCR, with a standard curve to ensure linear amplification. Immunoprecipitation efficiency was calculated relative to input = 1.

### Primers for ChIP

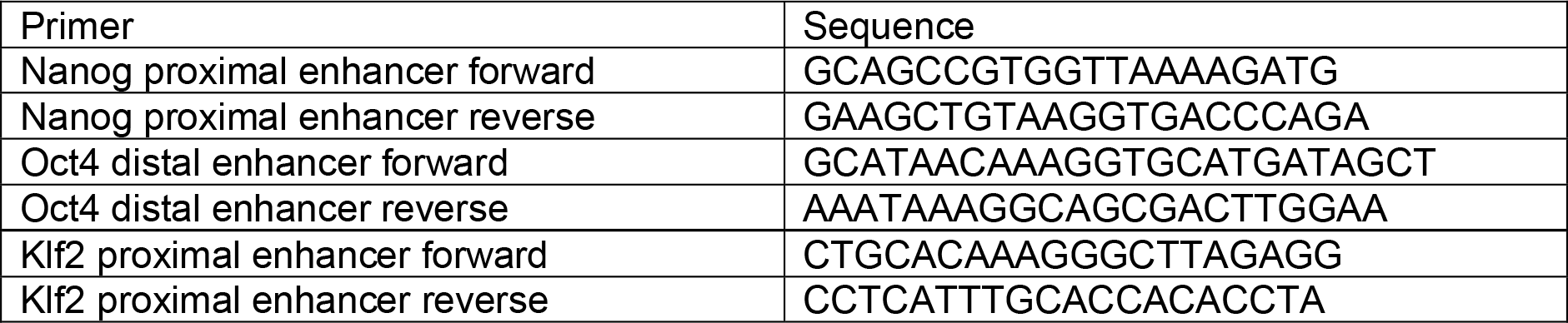

### RNA extraction, cDNA synthesis and qPCR

Total RNA was isolated using the RNeasy Mini kit (QIAGEN). 1µg RNA was reverse - transcribed using SuperScript III First-Strand Synthesis SuperMix for qRT-PCR (ThermoFisher Scientific). The resultant cDNA was analysed by quantitative PCR using TaqMan Fast Universal PCR Master Mix (ThermoFisher Scientific) with TaqMan Gene Expression Assays or with Fast SYBR Green Master Mix (ThermoFisher Scientific) using primers. qRT-PCR experiments were performed in triplicate on a StepOnePlus Real-Time PCR System (Applied Biosystems). Delta Ct values were normalised to GAPDH and raised to the power of −2. Standard deviations refer to technical replicates.

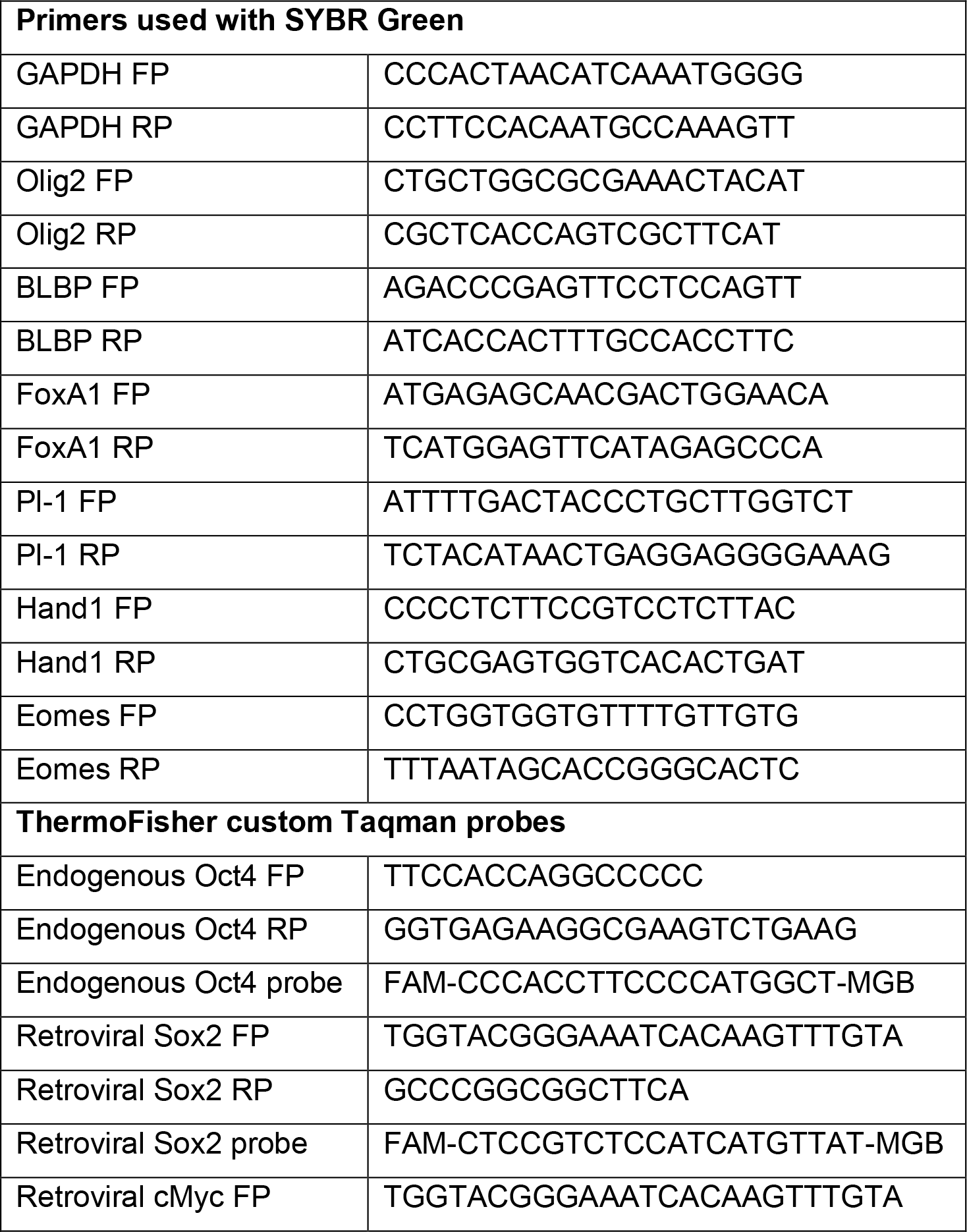
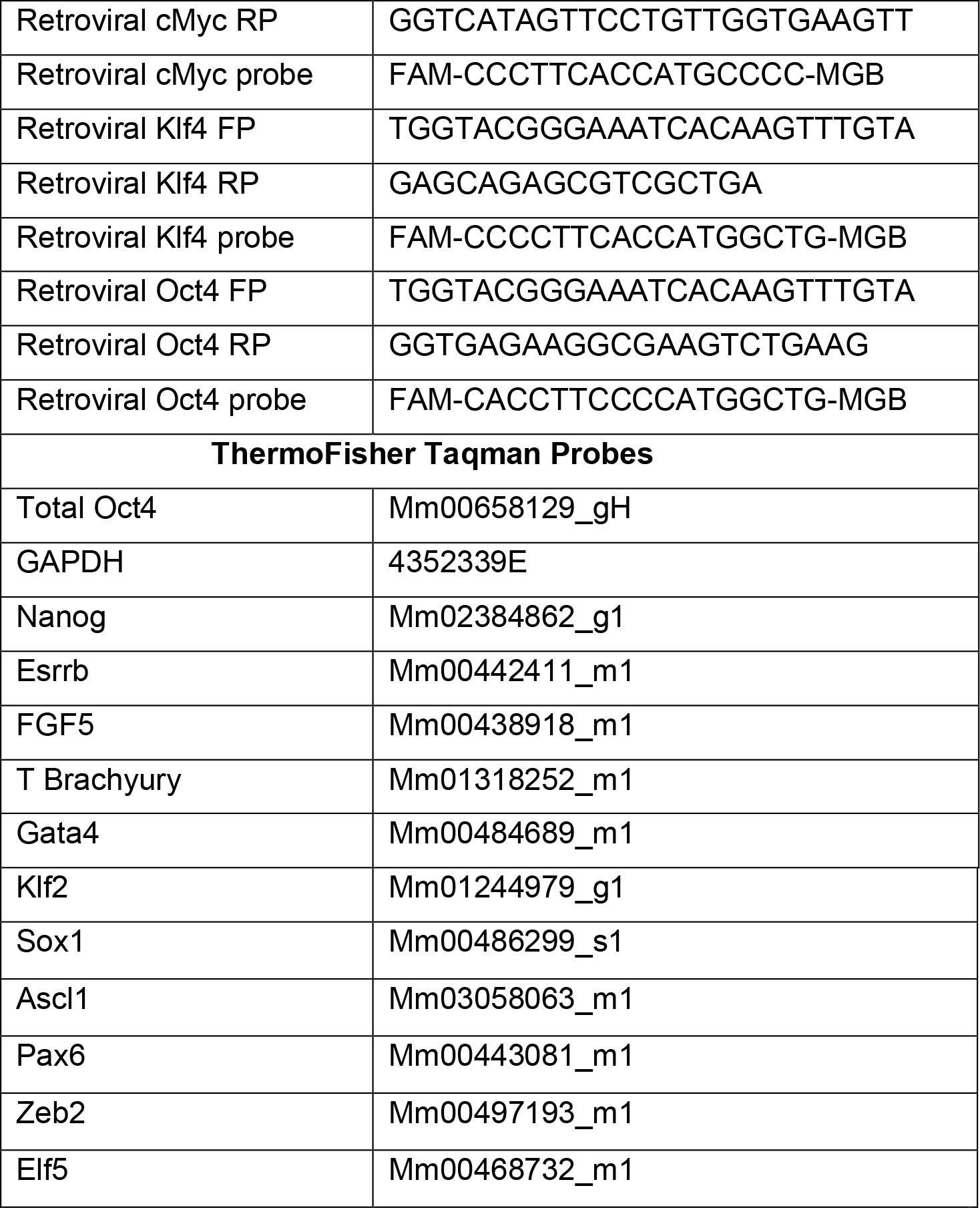

### scRNAseq library preparation

Single cells were index-sorted individually by FACS (BD Influx 5) into wells of a 96-well PCR plate containing lysis buffer. scRNAseq was performed as previously described (Nestorowa et al., 2016; Picelli et al., 2014; Wilson et al., 2015). The Illumina Nextera XT DNA kit was used to prepare libraries. Pooled libraries were sequenced on the Illumina HiSeq 4000 (single-end 125bp reads).

### RNAseq alignment

GENCODE M12 mouse gene annotation from Ensembl release 87 was used for read alignment (Yates et al., 2016) and splice junction donor/acceptor overlap settings were tailored to the read length of each dataset. Alignments to gene loci were quantified with HTSeq-count (Anders et al., 2015) based on annotation from Ensembl release 87. Quality control was performed according to (Stirparo et al., 2018). Briefly, sequencing libraries with fewer than 500,000 mapped reads were excluded from subsequent analyses. Read distribution bias across gene bodies was computed as the ratio between the total reads spanning the 50th to the 100th percentile of gene length, and those between the first and 49th. Samples with ratio >2 were not considered further. Stage-specific outliers were screened by principal component analysis.

### Published embryo scRNAseq datasets

Sequencing data of single-cell mouse embryo profiling studies SRP110669 (Mohammed et al., 2017) (E3.5, E4.5), SRP020490 (Deng et al., 2014) (trophectoderm cells) and E-MTAB-7901 (Stuart et al., 2019) (ESCs) were downloaded from the European Nucleotide Archive (Toribio et al., 2017) and from ArrayExpress repository and aligned as above.

### Transcriptome analysis

Principal component and cluster analyses were performed based on log2 FPKM values and were computed with FactoMineR (Lê et al., 2008) in addition to custom scripts. Default parameters were used unless otherwise indicated. For global analyses, genes that registered zero counts in all single-cell samples were omitted. Euclidean distance and complete linkage were used for cluster analyses unless otherwise indicated. Differential expression analysis was performed with scda (Kharchenko et al., 2014), that fits individual error models for assessment of differential expression between groups of cells. DAVID Bioinformatics Resources 6.7 (Huang et al., 2009) was used for computing the enriched biological processes, using as input list the modulated genes (with padj value < 0.05) between mutant cells and wt/rescue cells. Genes exhibiting the greatest expression variability (and thus contributing substantial discriminatory power) were identified by fitting a non-linear regression curve between average log2 FPKM and the square of coefficient of variation. Indicated specific thresholds were applied along the x-axis (average log2 FPKM) and y-axis (CV2) to identify the most variable genes.

